## Supplementary material for "Moving northwards: Invasive Green Crab (*Carcinus maenas*) Expands into the Southwestern Atlantic": Table S1. Complete dataset of bibliography review on green crabs population characteristics.: Supp_Mat_2_References_Table S1.docx

Aagaard, A.; Warman, C.G.; Depledge, M.H. Tidal and seasonal changes in the temporal and spatial distribution of foraging *Carcinus* *maenas* in the weakly tidal littoral zone of Kerteminde Fjord, Denmark. Mar. Ecol. Prog. Ser. 1995, 122, 165-172.

Acarli, S., Lök, A., Kirtik, A., Acarli, D., Serdar, S., Kucukdermenci, A., ... & Saltan, A. N. (2015). Seasonal variation in reproductive activity and biochemical composition of flat oyster (Ostrea edulis) in the Homa Lagoon, Izmir Bay, Turkey. *Scientia Marina*, *79*(4), 487-495.

Almeida, M. J., González-Gordillo, J. I., Flores, A. A., & Queiroga, H. (2011). Cannibalism, post-settlement growth rate and size refuge in a recruitment-limited population of the shore crab *Carcinus* *maenas*. *Journal of Experimental Marine Biology and Ecology*, *410*, 72-79.

Amaral, V., Cabral, H. N., Jenkins, S., Hawkins, S., & Paula, J. (2009). Comparing quality of estuarine and nearshore intertidal habitats for *Carcinus* *maenas*. *Estuarine, Coastal and Shelf Science*, *83*(2), 219-226.

Audet, D.; Miron, G.; Moriyasu, M. Biological Characteristics of a Newly Established Green Crab (*Carcinus* *maenas*) Population in the Southern Gulf of St. Lawrence, Canada. J. Shellfish Res. 2008, 27, 427-441.

Baeta, A.; Cabral, H.N.; Neto, J.M.; Marques, J.C.; Pardal, M.A. Biology, population dynamics and secondary production of the green crab *Carcinus* *maenas* (L.) in a temperate estuary. Estuar. Coast. Shelf Sci. 2005, 65, 43-52.

Baillie, C., & Grabowski, J. H. (2019). Invasion dynamics: Interactions between the European green crab *Carcinus* *maenas* and the Asian shore crab Hemigrapsus sanguineus. *Biological Invasions*, *21*(3), 787-802.

Béjaoui, B., Ismail, S. B., Othmani, A., Hamida, O. B. A. B. H., Chevalier, C., Feki-Sahnoun, W., ... & Hassen, M. B. (2019). Synthesis review of the Gulf of Gabes (eastern Mediterranean Sea, Tunisia): Morphological, climatic, physical oceanographic, biogeochemical and fisheries features. *Estuarine, Coastal and Shelf Science*, *219*, 395-408.

Bessa, F.; Baeta, A.; Martinho, F.; Marques, S.; Pardal, M.A. Seasonal and temporal variations in population dynamics of the *Carcinus* *maenas* (L.): The effect of an extreme drought event in a southern European estuary. J. Mar. Biol. Assoc. UK 2010, 90, 867-876.

Best, K.; McKenzie, C.; Couturier, C. Reproductive biology of an invasive population of European green crab, *Carcinus* *maenas*, in Placentia Bay, Newfoundland. Manag. Biol. Invasions 2017, 8, 247-255.

Chebbi, N., Mastrototaro, F., & Missaoui, H. (2010). Spatial distribution of ascidians in two Tunisian lagoons of the Mediterranean Sea. *Cahiers de Biologie Marine*, *51*(2), 117.

Dries, M.; Adelung, D. Die Schlei, ein Modell für die Verbreitung der Strandkrabbe *Carcinus* *maenas*. Helgol. Mar. Res. 1982, 35, 65-77.

Garbary, D. J., Miller, A. G., Williams, J., & Seymour, N. R. (2014). Drastic decline of an extensive eelgrass bed in Nova Scotia due to the activity of the invasive green crab (*Carcinus* *maenas*). *Marine biology*, *161*, 3-15.

Gillespie, G.E.; Phillips, A.C.; Paltzat, D.L.; Therriault, T.W. Status of the European Green Crab, *Carcinus* *maenas*, in British Columbia—2006. Canadian technical report of fisheries and aquatic sciences, Fisheries and Oceans Canada: Nanaimo, BC, Canada, 2007; p. 39.

Gillespie, G.E.; Norgard, T.C.; Anderson, E.D.; Haggarty, D.R.; Phillips, A.C. Distribution and Biological Characteristics of European Green Crab, *Carcinus* *maenas*, in British Columbia, 2006-2013. Canadian technical report of fisheries and aquatic sciences, Fisheries and Oceans Canada: Nanaimo, BC, Canada, 2015; p. 3120.

Himes, A. R., Balschi, W. S., Pelletier, G., & Frederich, M. (2017). Color phase—Specific ion regulation of the European green crab *Carcinus* *maenas* in an oscillating salinity environment. *Journal of Shellfish Research*, *36*(2), 465-479.

Hunter, E.; Naylor, E. Intertidal migration by the sore crab *Carcinus* *maenas*. Mar. Ecol. Prog. Ser. 1993, 101, 131-138.

Jensen, G. C., McDonald, P. S., & Armstrong, D. A. (2002). East meets west: competitive interactions between green crab *Carcinus* *maenas*, and native and introduced shore crab Hemigrapsus spp. *Marine Ecology Progress Series*, *225*, 251-262.

Jouili, S., Arculeo, M., Mansour, L., & Rabaoui, L. (2016). Biological characteristics of three Brachyuran crab species in the Lagoon of Elbibane, South-Eastern Tunisia. *Cah. Biol. Mar*, *57*, 217-226.

Kelley, A. L., de Rivera, C. E., Grosholz, E. D., Ruiz, G. M., Yamada, S. B., & Gillespie, G. (2015). Thermogeographic variation in body size of *Carcinus* *maenas*, the European green crab. Marine Biology, 162, 1625-1635.

Lyons, L.J.; O'riordan, R.M.; Cross, T.F.; Culloty, S.C. Reproductive biology of the shore crab *Carcinus* *maenas* (Decapoda, Portunidae): A macroscopic and histological view. Invertebr. Reprod. Dev. 2012, 56, 144-156.

Malvé, M. E., Battini, N., Livore, J. P., Schwindt, E., & Mendez, M. M. (2025). Spatial overlap and trophic interactions between a native commercial crab and the European green crab in Atlantic Patagonia. *Estuarine, Coastal and Shelf Science*, *312*, 109044.

McGaw, I.J.; Edgell, T.C.; Kaiser, M.J. Population demographics of native and newly invasive populations of the green crab *Carcinus* *maenas*. Mar. Ecol. Prog. Ser. 2011, 430, 235-240.

McVean, A. The incidence of autotomy in *Carcinus* *maenas* (L.). J. Exp. Mar. Biol. Ecol. 1976, 24, 177-187.

Monteiro, J. N., Ovelheiro, A., Ventaneira, A. M., Vieira, V., Teodósio, M. A., & Leitão, F. (2022). Variability in *Carcinus* *maenas* fecundity along lagoons and estuaries of the Portuguese coast. *Estuaries and Coasts*, 1-12.

Monteiro, J. N., Bueno-Pardo, J., Pinto, M., Pardal, M. A., Martinho, F., & Leitão, F. (2023). Implications of Warming on the Morphometric and Reproductive Traits of the Green Crab, *Carcinus* *maenas*. Fishes, 8(10), 485.

Monteiro, J. N., Ovelheiro, A., Maia, F., Teodósio, M. A., & Leitão, F. (2025). Biological traits and population dynamics for sustainable harvesting of *Carcinus* *maenas*. *Fisheries Research*, *281*, 107243.

Mouritsen, K.N.; Geyti, S.N.S.; Lützen, J.; Høeg, J.T.; Glenner, H. Population dynamics and development of the rhizocephalan Sacculina carcini, parasitic on the shore crab *Carcinus* *maenas*. Dis. Aquat. Org. 2018, 131, 199-211.

Nakano, H., Aikawa, T., Hagita, R., Hamada, H., Hayashi, T., Joshima, H., ... & Yoshida, J. (2023). Hydrographic structures of Tokyo Bay between 1992 and 2019 and evidence of temperature increase; observational results by the training vessel Seiyo-Maru. *Journal of Oceanography*, *79*(3), 281-294.

Naylor, E. Seasonal Changes in a Population of *Carcinus* *maenas* (L.) in the Littoral Zone. J. Anim. Ecol. 1962, 31, 601-610.

Odabaşı, S. Ü., Ceylan, Z., Şentürk, İ., Akbal, F., Bakan, G., & Büyükgüngör, H. (2022). Investigation of spatial and seasonal variation of water quality along the mid-Black Sea coast (from Sinop to Ordu) of Turkey, by multivariate statistical techniques. *Regional Studies in Marine Science*, *50*, 102169.

Quinn, B.K. Dramatic decline and limited recovery of a green crab (*Carcinus* *maenas*) population in the Minas Basin, Canada after the summer of 2013. PeerJ 2018, 6, e5566.

Reid, D.G.; Abello, P.; Warman, C.G.; Naylor, E. Size-related mating success in the shore crab *Carcinus* *maenas* (Crustacea: Brachyura). J. Zool. 1994, 232, 397-401.

Sheehan, E. V., Thompson, R. C., Coleman, R. A., & Attrill, M. J. (2008). Positive feedback fishery: population consequences of ‘crab-tiling’on the green crab *Carcinus* *maenas*. *Journal of Sea Research*, *60*(4), 303-309.

Souza, A.T.; Ilarri, M.I.; Campos, J.; Marques, J.C.; Martins, I. Differences in the neighborhood: Structural variations in the carapace of shore crabs *Carcinus* *maenas* (Decapoda: Portunidae). Estuar. Coast. Shelf Sci. 2011, 95, 424-430.

Tremblay, M.J.; Thompson, A.; Paul, K. Recent Trends in the Abundance of the Invasive Green Crab (*Carcinus* *maenas*) in Bras d'Or Lakes and Eastern Nova Scotia Based on Trap Surveys. Fisheries and Ocean Canada, Bedford Institute of Oceanography: Dartmouth, NS, Canada, 2006; p. 32.

Yamada, S.B.; Dumbauld, B.R.; Kalin, A.; Hunt, C.E.; Figlar-Barnes, R.; Randall, A. Growth and persistence of a recent invader *Carcinus* *maenas* in estuaries of the northeastern Pacific. Biol. Invasions 2005, 7, 309-321.

Yamada, S.B.; Gillespie, G.E. Will the European green crab (*Carcinus* *maenas*) persist in the Pacific Northwest? ICES J. Mar. Sci. 2008, 65, 725-729.

Young, A. M., & Elliott, J. A. (2020). *Life history and population dynamics of green crabs (Carcinus maenas). Fishes, 5 (1), 1–44*.
